# Enhanced simulations of whole-brain dynamics using hybrid resting-state structural connectomes

**DOI:** 10.1101/2023.02.16.528836

**Authors:** Thanos Manos, Sandra Diaz-Pier, Igor Fortel, Ira Driscoll, Liang Zhan, Alex Leow

## Abstract

The human brain, composed of billions of neurons and synaptic connections, is an intricate network coordinating a sophisticated balance of excitatory and inhibitory activity between brain regions. The dynamical balance between excitation and inhibition is vital for adjusting neural input/output relationships in cortical networks and regulating the dynamic range of their responses to stimuli. To infer this balance using connectomics, we recently introduced a computational framework based on the Ising model, first developed to explain phase transitions in ferromagnets, and proposed a novel hybrid resting-state structural connectome (rsSC). Here, we show that a generative model based on the Kuramoto phase oscillator can be used to simulate static and dynamic functional connectomes (FC) with rsSC as the coupling weight coefficients, such that the simulated FC well aligns with the observed FC when compared to that simulated with traditional structural connectome. Simulations were performed using the open source framework The Virtual Brain on High Performance Computing infrastructure.

## 1. Introduction

The human brain is a complex neural network that self-organises into different emergent states, crucial for its functions. Such states include spatiotemporal patterns of neural synchronisation associated with cognitive processes [1]. Brain regions can be modelled as dynamically interacting nodes in a functional network on a 3D space (functional brain networks), coupled in a complex manner driven by the structure of these networks. Over the past years, interdisciplinary approaches using concepts from nonlinear dynamics, physics, biology and medicine to name a few, allowed us to understand in more depth how the human brain functions, and how certain brain disorders and their underlying mechanisms can be further studied using mathematical models. It is feasible to ameliorate even more the predictive performance of such models, since a vast amount of neuroimaging data. e.g., electroencephalography (EEG), magnetoencephalography (MEG), and functional magnetic resonance blood-oxygen-level-dependent (BOLD) functional magnetic resonance imaging (fMRI) became available in the last 2 to 3 decades. Such data may provide information not only for healthy or pathological brain activity but can also be used to fingerprint functional connectomes by identifying individuals using brain connectivity patterns [2].

A dynamical-model approach can provide us with links between concepts from mathematics, physics, dynamical systems theory and empirically discovered phenomena. For example, it can provide us with links between attractors, bifurcations, synchronisation and empirical neuroimaging data. In this way, one can employ a mathematical model to replicate the time evolution of recordings from brain activity [3]. Hence, by choosing adequate model parameters, it is feasible to build customised virtual brain activity for individual subjects.

Together with extensive experimental work, mathematical/computational modelling of the whole brain dynamics has been an active research topic for years (e.g., [4, 5, 6, 7]). In such a setting one can model populations of neurons as nodes in a graph structure. Then, one can obtain information about relative connection weights (coupling strength) and communication lag (delay) between different nodes by diffusion-weighted magnetic resonance imaging (dwMRI) techniques (see e.g., [8, 9, 10]). This is termed as the structural connectivity (SC) of the network and it is in general subject-dependent with a certain degree of variability (gender/age/healthy vs diseased etc.). Furthermore, statistical analysis of BOLD time series inferred from fMRI can provide the functional relationships between different brain regions. It is usually calculated as the Pearson correlation coefficient of the activity between regions and results in the empirical functional connectivity (FC) matrix per brain recording and subject (see e.g., [11, 12]).

By working on a “virtual” environment, one can seek for model parameters that are able to produce simulated time series and global dynamics that fairly resemble the empirical ones. One way in achieving that is to tune selected parameters which optimize the similarity between empirical FC with the simulated FC (see e.g., [13, 14]). Hence, these parameters can serve as dynamical biomarkers and predictors of different brain states and behavioural modes (see [15] for a recent review). Along this direction, the virtual epileptic patient has been recently proposed, where medical-treatment approaches using personalised mathematical models for epileptic patients have been illustrated (see e.g., [3]). Furthermore, the choice of the brain atlases, (i.e. the mapping of the different regions of interest (ROIs) based on functional or anatomical criteria using different parcellations) can affect the quality of model performance and its level of agreement with the empirical data (see [16] and references therein for more details).

In recent years, substantial research efforts have been directed toward understanding the brain (large-scale activity) using resting state fMRI (rs-fMRI) employing sophisticated mathematical and statistical tools to investigate the FC from rs-fMRI data [17]. So far, the mainstream approach is to consider SC to be static and the FC one dynamic. However, this is not necessarily the case as white matter tracts can be in use or engaged when the brain is performing certain tasks but inactive or disengaged during other tasks and hence not static. An altered and more sophisticated “functional connectivity-informed structural connectivity” has been introduced in [18] employing information from fMRI to infer the underlying pattern of white matter engagement specific to the brain’s state. The resulting connectome, the so-called resting-state informed structural connectome (rsSC), encodes the structural network that underlies and facilitates the observed rs-fMRI correlation connectome able to detect altered rsSC community structure in diseased subjects relative to controls. In the original set up there is no “directionality” inferred, i.e., whether the white matter tract of interest is of “excitatory” versus “inhibitory” nature.

However, understanding the dynamical balance between excitation and inhibition, a concept termed E-I balance, is vital for adjusting neural input/output relationships in cortical networks and regulating the dynamic range of their responses to stimuli [19] such that information capacity and transfer are maximized [20]. This is the central thesis of the criticality hypothesis [21], i.e., that brain activity self-organize into a critical state [22], a unique configuration likened to a phase transition in physical systems where a dynamical system transitions from order (balanced excitation-inhibition) to disorder (disrupted excitation-inhibition balance) [23, 24, 25, 26]. Indeed, evidence supporting that the brain is operating near criticality has been reported in studies examining neuronal signaling [24, 27, 28] as well as BOLD fMRI signals [29, 30, 31, 32]. Optimal E-I balance is crucial for healthy cerebral activity and maintaining homeostatic control, while dysfunction of this balance may lead to neurological disorders. For example, autism has been shown to exhibit a shift towards a more excitatory state [33, 34], while Down syndrome has been linked to an increase in inhibition (or decrease in excitation) [35]. Additionally, there is growing evidence suggesting that neuronal hyper-excitation may represent the earliest changes in AD due to the susceptibility of inhibitory GABAergic interneurons to apolipoprotein E (APOE) *ε*4 -mediated neurotoxicity (APOE *ε*4 is the strongest genetic risk factor for AD) [36, 37, 38, 39].

To incorporate co-activation (excitatory) or silencing (inhibitory) effects into our hybrid rsSC framework that would allow us to infer the brain’s E-I balance, in [40] we then introduced an improved framework based on the Ising model representation of the brain as a dynamical system, wherein self-organized patterns are formed through the spontaneous fluctuations of random spins. This Ising spin-glass model has been previously used to successfully characterize complex microscale dynamics [4, 41, 42, 43] and macroscale interactions [44, 45, 46, 47, 48, 49] of the human brain, and to accurately represent spatiotemporal co-activations in neuronal spike trains [49, 50, 51] and patterns of BOLD activity [52, 53, 54, 55]. Using this improved “signed” rsSC (where the magnitude of the rsSC edge weights is constrained by the structural connectivity while the sign of the edges indicates either excitatory or inhibitory), we were able to demonstrate female-specific age-associated hyper-excitation in a group of APOE *ε*4 cognitively-normal women carriers, suggesting our approach is a feasible path towards an E-I balance based imaging biomarker for AD.

In this paper, we use The Virtual Brain (TVB, [5]), a whole-brain simulation platform part of the EBRAINS infrastructure [56], to investigate the potential benefits in employing rsSC instead of the traditional SC for simulating whole-brain dynamical activity. For example, one major limitation when employing certain dynamical models, such as the Kuramoto phase oscillators [57] and the generic limit-cycle oscillators [58], to model each node’s mean neural activity using traditional SC connectomes is that the resulting signals from different ROIs do not produce negative correlations similar to the empirical ones, even in the presence of delays in the system (see e.g. [16]). However, this is not the case in time series obtained by empirical neuroimaging data where both positive and negative correlations coexist. We here show that by using rsSC such dynamical systems are able to produce simulated signals with both positive and negative correlations following the trends of the empirical ones. Furthermore, we show that in general the agreement between simulated FC and empirical FC matrices (representing the respective BOLD signal activity) is improved significantly when rsSC is used instead of standard SC.

## 2. Methods and materials

### 2.1. Empirical data

Structural and functional connectivity for 38 cognitively normal APOE *ε*4 allele carriers aged 40–60 (*μ* = 50.8) are compared with 38 age (*μ* = 50.9) and sex-matched (16 male/22 female) non-carriers (control - non-carriers). Resting state functional MRI (rs-fMRI)—A T2*-weighted functional scan was obtained with an echo-planar pulse imaging (EPI) sequence (28 axial slices, 20 × 20 cm^2^ FOV, 64 × 64 matrix, 3.125 mm × 3.125 mm × 4 mm voxels, TE = 40 ms, TR = 2,000 ms). The 8-minute rs-fMRI scan was acquired under a task-free condition (i.e., resting state): subjects were instructed to relax with eyes closed and to “not think about anything in particular”. Imaging included T1-weighted MRI, resting state fMRI and diffusion weighted MRI. Freesurfer cortical parcellation and sub-cortical segmentation was performed to derive 80 regions-of-interest (ROIs) registered on the Desikan atlas [59]. The mean time-course was extracted from the pre-processed rs-fMRI data. Probabilistic tractography was used to create the structural connectome matrices, and normalized by the way-total of the corresponding seed ROIs. The detailed information on the imaging and processing steps can be found in [60].

### 2.2. Signed resting state structural connectome

In constructing a signed resting state structural connectome, we use a novel approach introduced in [18] and has already been used in several studies (see e.g. [40, 61, 21] which takes into account both structural connectivity and functional time series to form a signed coupling interaction network or “signed resting state structural connectome” (signed rsSC) to describe neural excitation and inhibition. To this end, an energy representation of neural activity based on the Ising model from statistical mechanics which ultimately bypasses traditional BOLD correlations. The spin model is a function of a coupling interaction (with positive or negative values) and spin-states of paired brain regions. Observed functional time series represent brain states over time. A maximum pseudolikelihood with a constraint is used to estimate the coupling interaction. The constraint is introduced as a penalty function such that the learned interactions are scaled relative to structural connectivity; the sign of the interactions may infer inhibition or excitation over an underlying structure. The efficiency of this approach was validated in comparing a group of healthy APOE *ε*4 carriers (associated with genetic risk factor for Alzheimer’s disease with a control (healthy) group of non APOE-*ε*4 subjects.

Here, we briefly describe the computational aspect of this approach. First, we adapted the Ising model, a well-known spin-glass model from statistical physics in which the states, also referred to as “spin configurations”, of interacting units – in our case brain regions connected by white matter edges – are constrained to be either 1 (“active”) or -1 (“inactive”). As described in [21, 62] we construct a function-by-structure embedding (FSE) using a constrained pseudolikelihood estimation technique wherein pairwise interaction coefficients (represented as (*J*_*i,j*_), with *i* and *j* representing ROIs in the brain network) are inferred from the cobserved data (BOLD time series). As the model assumes binary data, we binarize the resting-state fMRI signals. The binarized activity pattern of all ROIs at time *t* (*t* = 1, 2, …, *t*_max_) is (**s**(*t*) = *s*_1_(*t*), *s*_2_(*t*), … *s*_*N*_ (*t*) ∈ {−1, +1}^*N*^).

Note that *t*_max_ is determined as a result of the fMRI scan time. Here (*s*_1_(*t*) = ±1) indicates that an ROI is either active (+1) or inactive (−1). First, the time series goes through a z-score normalization procedure, resulting in zero mean and unitary variance. The interaction (*J*_*i,j*_) between two regions should be directly linked back to the diffusion MRI-derived structural connectivity between them as informed by tractography, so we add a constraint to the Hamiltonian function as:

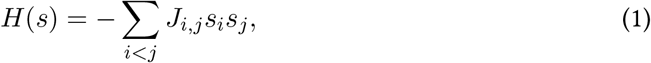

such that |*J*_*i,j*_| ∝ *W*_*i,j*_, where (*W*_*i,j*_) is the structural connectivity between pairs of ROIs, and the external force or bias terms are dropped in the case of resting-state. This ensures that in the pseudolikelihood estimation of (**J**), we constrain it with the structural connectivity (under the assumption that structural connectivity informs spin models governing brain dynamics). Thus, the optimal interaction matrix (**J**) is derived by maximizing the pseudo-likelihood function as:

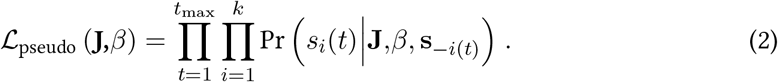

Pseudolikelihood substitutes Pr(*s*(*t*) by the product of the conditional probabilities 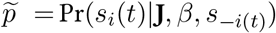 observing one element *s*_*i*_(*t*) with all the other elements (denoted *s*_−*i*(*t*)_) fixed. To ensure that the magnitude of the coupling interactions is scaled relative to structural connectivity, the constraint is formulated as |*J*_*i,j*_| ≈ *μW*_*i,j*_, where *μ* is a normalization constant and *W*_*i,j*_ is the structural connectivity between ROI pairs. Without loss of generality, we assume that *μ* = 1 with appropriate normalization. We therefore present a penalty-based optimization scheme to maximize the constrained log-pseudolikelihood function as:

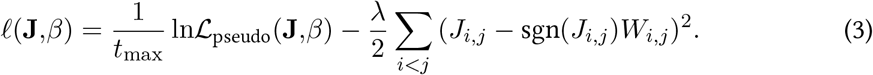

And the pseudolikelihood component expands as follows:

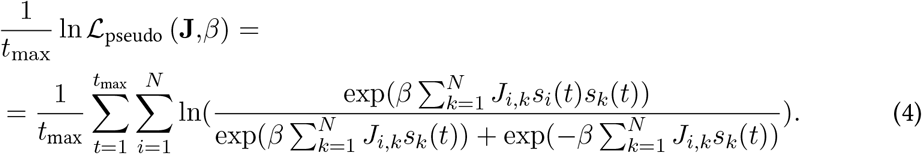

Our formulation here is based on the Boltzmann distribution under pseudolikelihood conditions. Thus, the numerator describes the energy of the system, while the denominator is the sum of all possible energies. Hence, there are only two terms in the denominator since *s*_*i*_(*t*) is binary (one positive, and one negative). The likelihood function may be simplified by setting 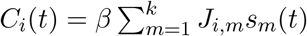, resulting in:

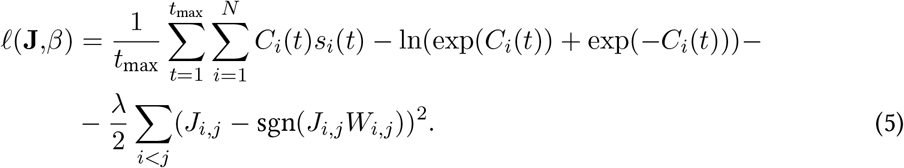

Here we may construct the gradient ascent procedure with respect to *J*_*i,j*_ by computing the partial derivative of the log-pseudolikelihood as:

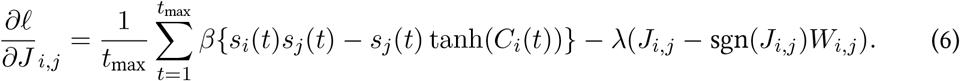

The updating scheme follows:

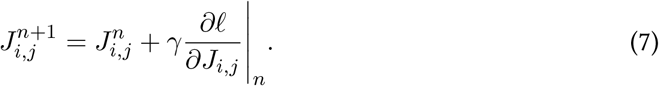

Here, *n* is the iteration number and *γ* is the learning rate. In this way, the penalty function ensures that the inferred pairwise interaction is scaled relative to the estimated structure of the brain. This procedure is followed for all subjects in constructing an optimized **J** matrix per subject, which we term the resting-state structural connectome or rsSC.

### 2.3. Models and simulated data

In order to produce simulated fMRI time series in the given connectomes, we employ the Kuramoto phase oscillator model [57, 63, 16]:

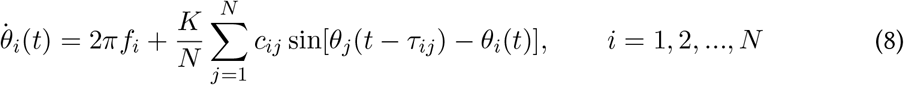

where *θ*_*i*_ are the phases, *N* is the number of oscillators, *f*_*j*_ are the natural frequencies (Hz), *c*_*ij*_ and *τ*_*ij*_ (ms) represent the individual coupling weight and propagation delay in the coupling, respectively, from oscillator *j* to oscillator *i* while *K* is the global coupling parameter. The time *t* in the model and delay in coupling term are measured in ms.

For each individual subject, we produced a “personalized” model Eq. (8) to simulate the network’s dynamics and to calculate time series. To this end, two cases of connectivity matrices were compared: (i) in the first one the *c*_*ij*_ values are defined by simply counting the number of streamlines connecting regions *i* and *j* normalized to 1 and with zero diagonal (i.e., define *c*_*ij*_ as a normalized version of the empirical tractography-derived SC or eSC), leading to only excitatory interactions between ROIs; and (ii) in the second one the *c*_*ij*_ values are assigned by the corresponding entries of the hybrid rsSC connectomes, leading to both excitatory and inhibitory interactions between ROIs. The delays *τ*_*ij*_ were calculated as *τ*_*ij*_ = *L*_*ij*_/*V*, where *L*_*ij*_ (mm) is the average tract (path) length of the streamlines connecting regions *i* and *j*, and *V* (m/s) is an average velocity of signal propagation. In this particular dataset the exact path lengths are not available, hence we used instead the euclidean distance between nodes as proxies. The euclidean distance has been used in the literature in the construction of structural networks (see e.g. [64]) and found to closely follow the trends obtained by anatomical tract-tracing studies. Furthermore in [65], the authors showed that such networks also strongly correlate with MRI tractography-based networks. The matrix **L** = *L*_*ij*_ can thus be used to calculate the delays *τ*_*ij*_ in the coupling, which can be expressed as *τ*_*ij*_ = *τ* · *L*_*ij*_/⟨*L*_*ij*_⟩, where *τ* = ⟨*L*_*ij*_⟩/*V* is the global (or mean) delay. In Eq. (8) the self-connections were excluded by setting the diagonal elements in the matrices eSC/rsSC and **L** to zero (i.e. *c*_*ii*_ = *L*_*ii*_ = 0 respectively).

In **Figure 1**(A), we show the empirical SC matrix (weights of the node-to- node connections) for a Non Carrier subject from the dataset with 80 nodes (ROIs). **Figure 1**(B) depicts the corresponding to this SC and subject rsSC matrix calculated as described earlier. Note that the hybrid rsSC contains negative entry values as opposed to SC one that is restricted to having only positive values. **Figure 1**(C) shows the tract length **L** matrix (in mm) that we used for all subjects’ simulation in the absence of the actual measured ones from a neuroimaging prepossessing pipeline.

**Figure 1:**
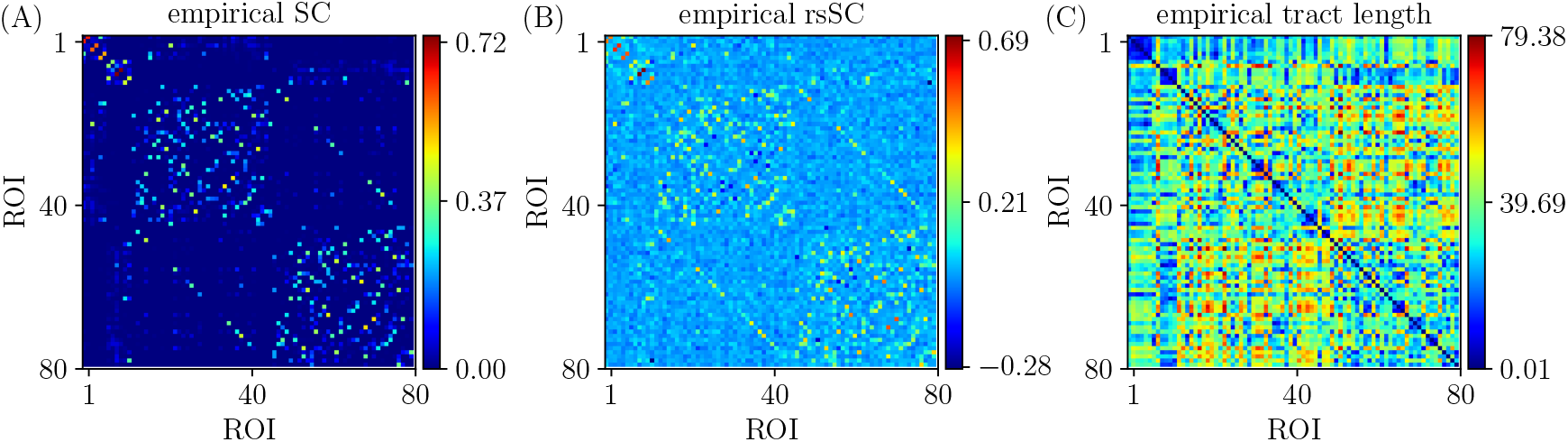
Empirical connectivity matrices example (non-carrier subject). (A) Weights of SC matrix. (B) Hybrid rsSC matrix. (C) Tract length (mm) matrix **L** based on the euclidean distance of the nodes on the Desikan atlas (same for all simulations and subjects).

The phases in our model (Eq. 8) were initialized randomly. We set the intrinsic frequencies to be uniformly distributed with mean = 60 Hz and SD = 1 Hz, corresponding to oscillations within the gamma frequency range (see e.g. [13, 66, 67, 63] for more details and motivation), as gamma local field potential (LFP) power is coupled to the BOLD fMRI signal and is considered representative of the overall neuronal activity (see also [68, 69, 70, 71]).

For our simulations, we used a TVB tailored version for the Kuramoto model and we made adjustments for efficient parallelisation on CPUs using MPI on the supercomputer JUSUF located at the Jülich Supercomputing Centre. Our model generates time series which correspond initially to electrical activity (fast oscillations) for each node, i.e. we register the observable *x*_*i*_ = sin(*θ*_*i*_) for each brain region. Then, in order to estimate the simulated BOLD signal, we use the TVB’s build-in tool to calculate the haemodynamic response function kernel (i.e. “fMRI activity”) associated with a given neural activity time series, also known as the Balloon-Windkessel model [72]. Our simulations ran for 500 seconds in total. The first 20 seconds were discarded to remove transient effects, resulting in *T* = 480 seconds (8 minutes), i.e. a time interval identical to the time-length of the empirical fMRI signals. We set the time-step at 0.1 ms and we integrated the system with an Euler scheme. In this particular study we did not consider the presence of noise.

## 3. Results

We numerically simulate BOLD (ultimately) time series varying two model parameters, namely the global coupling strength *K* and the delay *τ* in Eq. (8), with respective ranges *K* ∈ [1, 75] and *τ* ∈ [1, 33] resulting in a 32 × 32 grid. For each pair of parameters, we begin by producing the matrix of the simulated FC (sFC). The latter is measured by the Pearson Correlation Coefficient (CC) between the simulated BOLD signals *x*_*i*_, *i* = 1, 2, …, *N* from different ROIs (also referred to as Static Functional Connectivity in the literature, see e.g. [73]), namely:

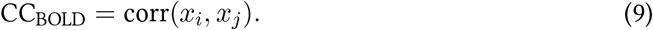

Then, we compare each sFC with the eFC ones using, again, the Pearson Correlation Coefficient, however this time we calculate it for the two respective matrices (upper triangular parts), i.e.:

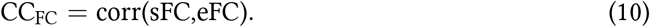

The optimal match between sFC and eFC in the parameter space is acquired for (*K, τ*)-values where CC_FC_ becomes maximal (see also [16] and references therein for more details and motivation).

In **Figure 2**, we present the first main result, namely, the superiority of hybrid rsSC over standard SC matrices in generating simulated BOLD time series with models like Eq. (8) which better approximate the empirical BOLD signals (shown here for one example healthy subject). The upper row refers to simulations performed using the respective subject’s standard SC matrix to define the coupling weights in Kuramoto model. **Figure 2**(A) shows the parameter sweep exploration (PSE) for eFC vs. sFC and for the parameters (*K, τ*) and measured as CC_FC_ = corr(sFC,eFC). The 5 white circles on the red regions indicate the highest correlations found (larger circles’ sizes correspond to larger CC_FC_ values). In **Figure 2**(B) we present the eFC calculated from the empirical BOLD signal while in **Figure 2**(C) the sFC matrix with the larger cc_FC_. We can observe that sFC did not capture adequately the negative correlations that are present in eFC (compare the minimum values in the barplots of panel (B) and (C)).

**Figure 2:**
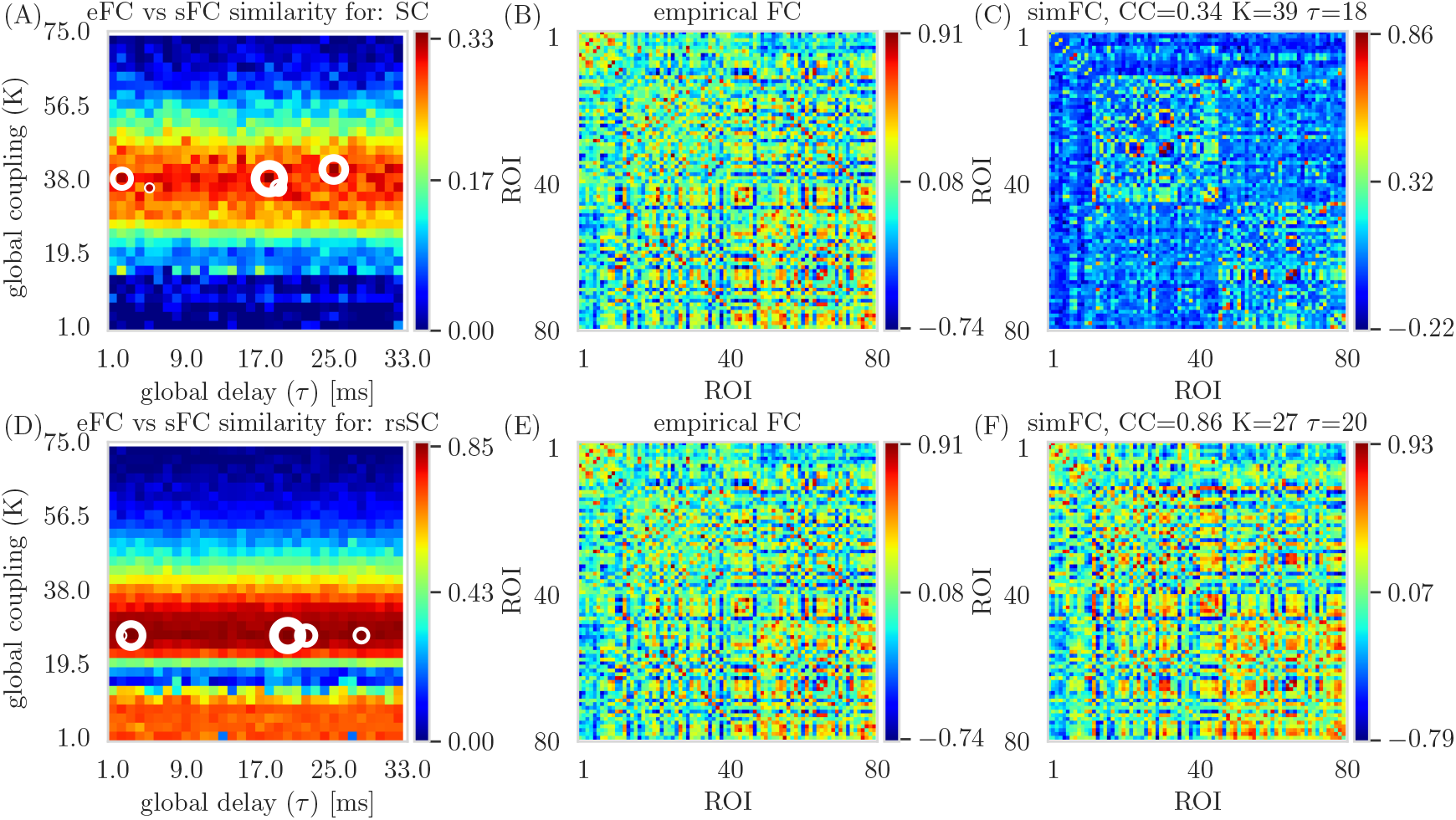
Parameter Sweep Exploration for eFC vs. sFC example. Upper row (using the respective subject’s standard SC matrix to define the weights in model Eq. (8): (A) The colormap depicts the CC_FC_ = corr(sFC,eFC) for the parameters (*K, τ*). The 5 circles on the red regions indicate the highest correlations found (larger circles’ sizes correspond to larger CC_FC_ values). (B) The eFC calculated from the empirical BOLD signal. (C) The sFC matrix with the larger CC_FC_. Lower row (using the respective subject’s hybrid rsSC matrix to define the weights in model Eq. (8): (D) The respective colormap for CC_FC_ = corr(sFC,eFC). (E) The eFC calculated from the empirical BOLD signal (same as (B)). (E) The sFC matrix with the larger CC_FC_. Note the ranges for the two colorbars in (A) and (D) are kept intact (i.e. no scaling) for visualization purposes.

In the lower row of **Figure 2** (panels (D),(E),(F)), we perform similar simulations, however we now use the respective subject’s hybrid rsSC matrix to define the coupling weights in the Kuramoto model). Note the significant improvement in the maximum value of the CC_FC_ ≈ 0.86 compared to the one found when using the standard SC matrix (CC_FC_ ≈ 0.33). Note also the better agreement between the two FC matrices (empirical (D) and simulated (E)) and how better the sFC captures both positive and negative correlations (indicated by the range of the respective colorbars). We should stress that we did not opt to use the same range for the two colorbars in **Figure 2**(A) and (D), as in this way it would be difficult to visually identify the PSE region in (A) depicting the optimal parameter values.

The respective scatterplots and CC values between empirical and optimal simulated FC matrices are presented in **Figure 3** using rsSC (A) and standard SC (B) matrices. Here we plot the empirical (y-axis) against the optimal simulated BOLD correlations (x-axis) aggregated across all entries in the corresponding FC matrices, thus a perfect match between the two would place all the points along the line x=y. The higher CC_FC_(sFC,eFC) value (using hybrid rsSC matrices) is well reflected by a rather clear linear trend in the distribution of the points (panel (A)), On the other hand, only a relatively weak linear trend is obtained using standard SC matrices (panel (B)). Both panels refer to same subject presented in **Figure 2** with the lines indicating the corresponding linear fit in each case.

**Figure 3:**
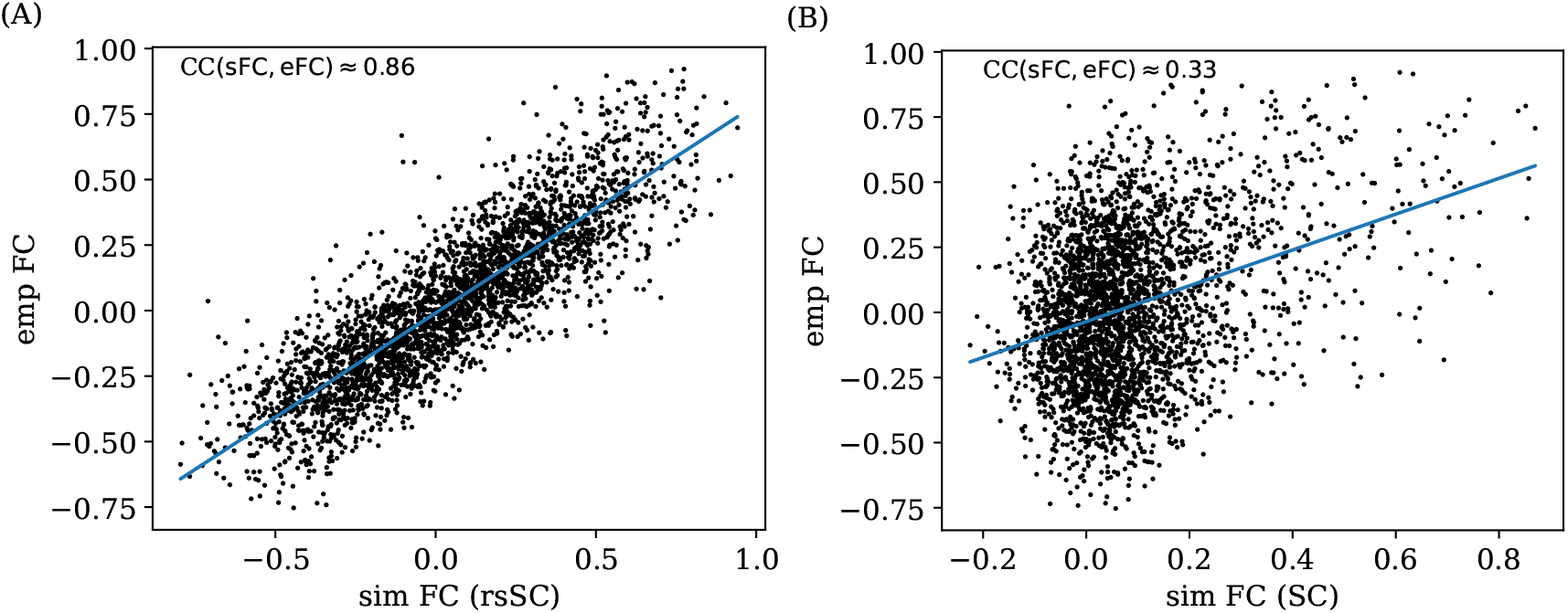
Correlation Analysis. Scatterplots between empirical (*y*−axis) and optimal simulated BOLD correlations (*x*−axis) aggregated across all entries in the corresponding FC matrices, i.e., panels (B) and (C) in Figure 2. Note that a perfect match between the two FC matrices would thus place all the points along the line *x* = *y*. (A) eFC vs. sFC using rsSC (B) eFC vs. sFC using standard SC. Both panels refer to same subject presented in **Figure 2**). The blue lines indicate the respective linear regression model.

In **Figure 4** we present a statistical analysis for all subjects per category, i.e. 38 non-carriers (A) and 38 carriers (B). For each subject we considered the 5 maximum values of correlation coefficients between eFC and sFC using SC and rsSC matrices for the simulated time series respectively (circles in **Figures 2**(A),(D)) and produced boxplots. We then used the t-test to compare the respective mean values. The difference in the respective mean values of the two datasets is found to be statistically significant with very small *p*−value (*p* ≤ 0.0001) for both non-carriers (**Figure 4**(A)) and carriers sets (**Figure 4**(B)).

**Figure 4:**
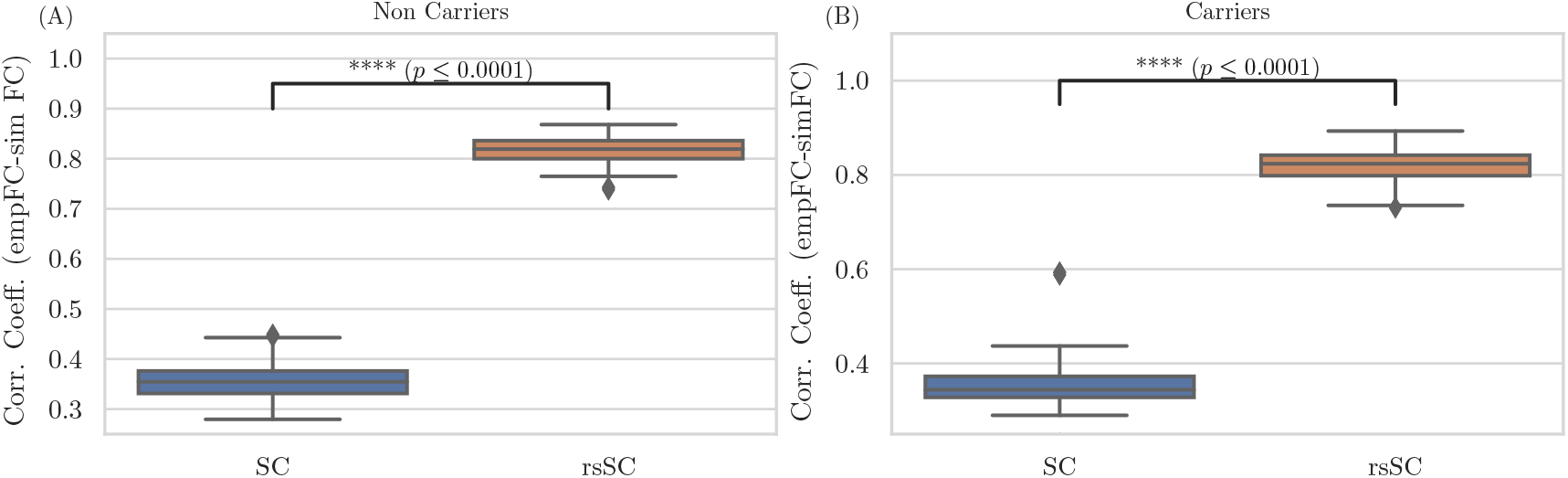
Statistical Analysis. Boxplots for the non-carriers (A) and carriers (B) dataset (38 subjects per type) correlation coefficients between eFC and sFC using SC and rsSC matrices for the simulated time series respectively. For each subject we considered the 5 maximum values (see circles in **Figures2**(A),(D)). The difference in the respective mean values of the two datasets is statistically significant measured by the t-test with very small *p*−value (*p* ≤ 0.0001) for both non-carriers and carriers sets.

In **Figure 5** we show the respective BOLD time series, In more detail, in **Figure 5**(A) we show the time evolution of the empirical BOLD signal. **Figure 5**(B) depicts the simulated BOLD signal with the parameters (*K, τ*) ≈ (27, 20) found when optimizing CC_FC_ using the respective rsSC matrix while in **Figure 5**(C) we plot the respective simulated BOLD signal with the parameters (*K, τ*) ≈ (39, 18) using the SC matrix. All BOLD signals in all three panels are scaled to range in [−1, 1].

**Figure 5:**
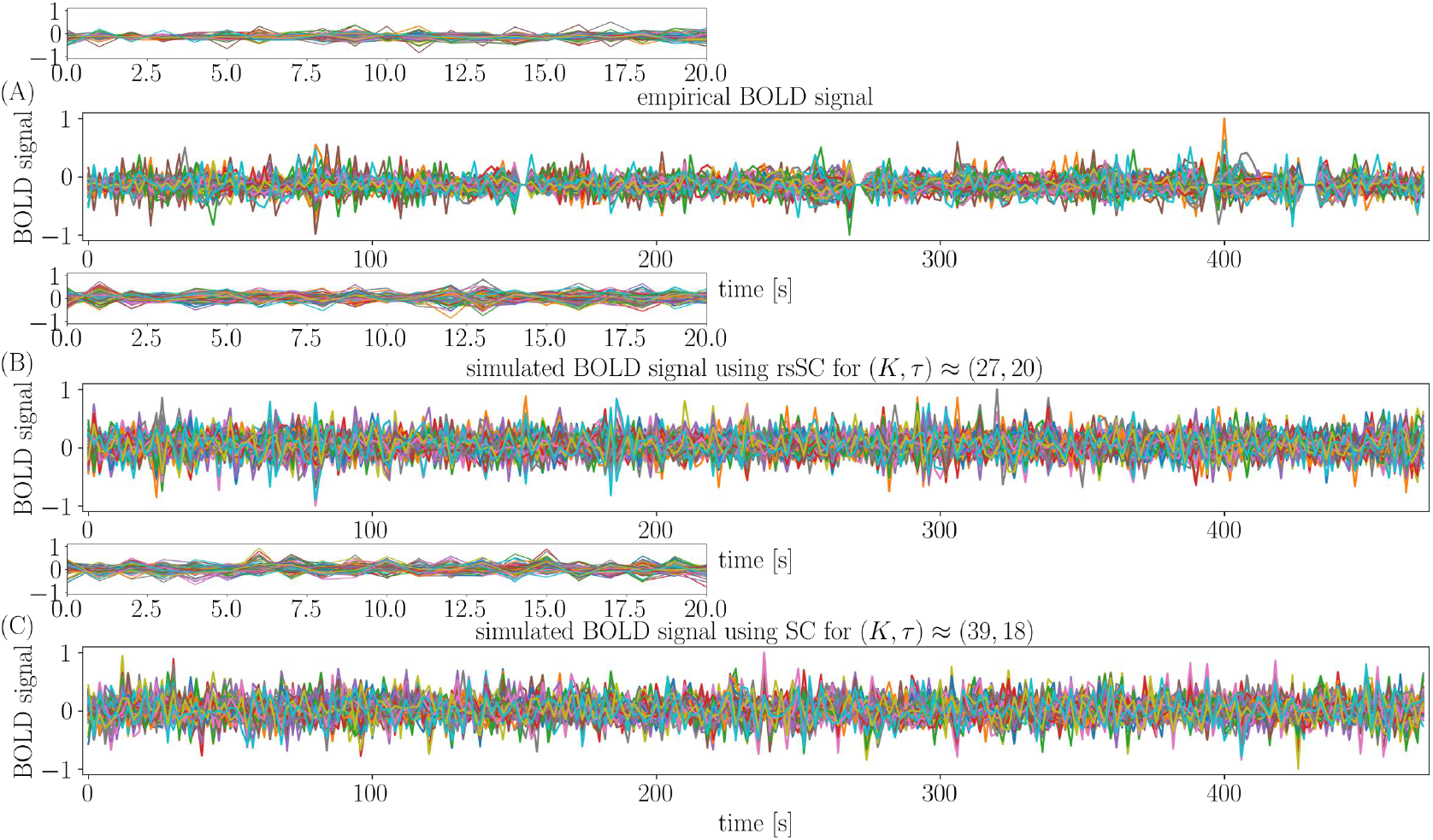
Bold signals. (A) Empirical BOLD signals. (B) Simulated BOLD signals with the parameters (*K, τ*) found when optimizing CC_FC_ using the respective rsSC matrix. (C) Simulated BOLD signals with the parameters (*K, τ*) found as in (B) but using the respective SC matrix. Note that all BOLD signals in all three panels are scaled to the range [−1, 1]. The small figures on top of the main panels show the respective zoomed areas for the first 20 seconds.

In order to quantify the better performance of the simulated BOLD signals generated when using the rsSC matrices, we calculated the minimum and maximum Pearson correlation coefficients CC_**BOLD**_ (Eq, (9)) in eFC and sFC (for rsSC and SC) matrices for all subjects (Non Carriers and Carriers separately). To this end, we considered the sFC produced by the 5 optimal (*K, τ*) parameters when fitting for eFC (see the 5 circles in **Figure 2**(A),(D) where an example for one subject is shown). In **Figure 6**, we show boxplots for the minimum/maximum CC_**BOLD**_ values for the eFC (light green) vs sSF using the rsSC (light blue) and SC (red) matrices (see e.g. minimum/maximum values of the colorbars of panels **Figure 2**(B),(C),(E),(F)) for Non Carriers (A) and Carries (B) respectively.

**Figure 6:**
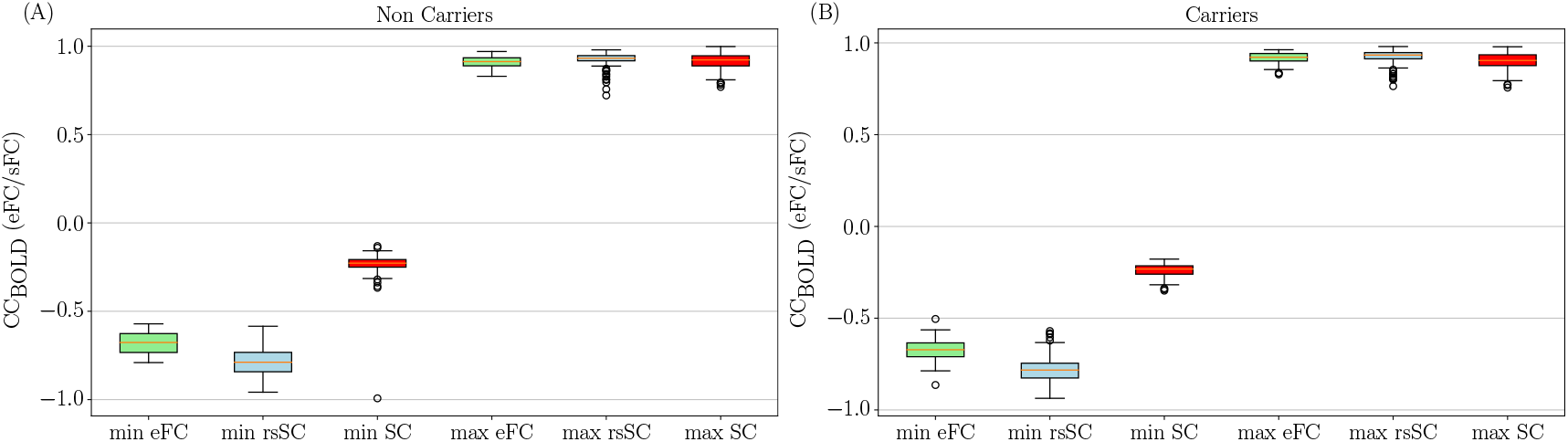
BOLD signal minimum and maximum Pearson correlation coefficients. (A) Boxplots of the minimum and maximum CC_**BOLD**_ values (Eq, (9)) for Non Carriers. These values are obtained by the sFC matrices produced by the 5 optimal (*K, τ*) parameters when fitting eFC (light green) with sFC using the rsSC (light blue) and SC (red) matrices (see **Figure 2**(A),(D)) and the minimum/maximum values of the colorbars of panels **Figure 2**(B),(C),(E),(F). The simulated time series obtained with SC (left red boxplots) do not succeed to capture adequately the negative correlations that occur in the empirical time series (left light green boxplots). Simulated signals generated with rsSC (left light blue boxplots) acquire negative correlations much closer to the empirical ones. (B) Similar analysis as in panel (A) for Carriers.

Both sFC (simulated with rsSC/SC) perform rather well in capturing the positive correlations observed in the empirical BOLD signals. However, note that the simulated time series obtained with SC (left red boxplots) do not adequately capture the negative correlations that occur in the empirical time series (left light green boxplots). On the other hand, BOLD signals generated with rsSC (left light blue boxplots) yield negative correlations much closer to the empirical ones. This trend is present for both Non Carriers and Carriers datasets.

Next, to further explore the advantage in using the hybrid rsSC matrices beyond Static Functional Connectivity matrices, we sought out to perform a similar PSE analysis for Dynamic Functional Connectivity, which allows us to capture switching trends in the resting-state activity. To this end, we calculate the Phase Coherence Connectivity (see e.g. (see e.g. [73, 74] and references therein) which does not suffer from time-window length effects like other similar techniques based on calculating successive FC(*t*) matrices using a sliding-window (see discussion in [73, 74]). Hence, we use BOLD Phase Coherence Connectivity to measure time-resolved dynamic FC matrices (dFC), with size *N* × *N* × *T*, where *N* refers to the number of ROIs and *T* = 236 the total number of recording frames. Then, we begin by estimating the phases from the BOLD time series (empirical and simulated) for all ROIs *i* (*θ*(*i, t*)) applying a Hilbert transform and we bandpass filter the parcellated fMRI time-series within 0.01 − 0.1 Hz (see e.g. [16] and references therein) using a discrete Fourier transform computed with a fast Fourier transform. Then, the phase coherence between brain areas *i* and *j* at time *t*, dFC(*i, j, t*) is defined as:

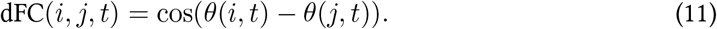

When two ROIs have temporarily aligned BOLD signals their respective dFC(*i, j, t*) ≈ 1 while their BOLD signals are orthogonal dFC(*i, j, t*) ≈ 0. Note the matrix dFC serves as the foundation of Leading Eigenvector Dynamic Analysis (LEiDA) which has been used to detect subtle FC patterns that distinguish healthy versus diseased BOLD signals (see e.g. [73, 74]).

In **Figure** 7, we present the Phase Coherence Connectivity PSE for edFC vs. sdFC (in a similar way as in **Figure** 2). However, we now compare each simulated mean dFC (sdFC) calculated by Eq. (12) with the empirical (edFC) ones using the Pearson Correlation Coefficient, from the upper triangular section of the two respective matrices, i.e.:

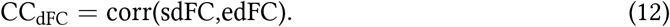

**Figure 7:**
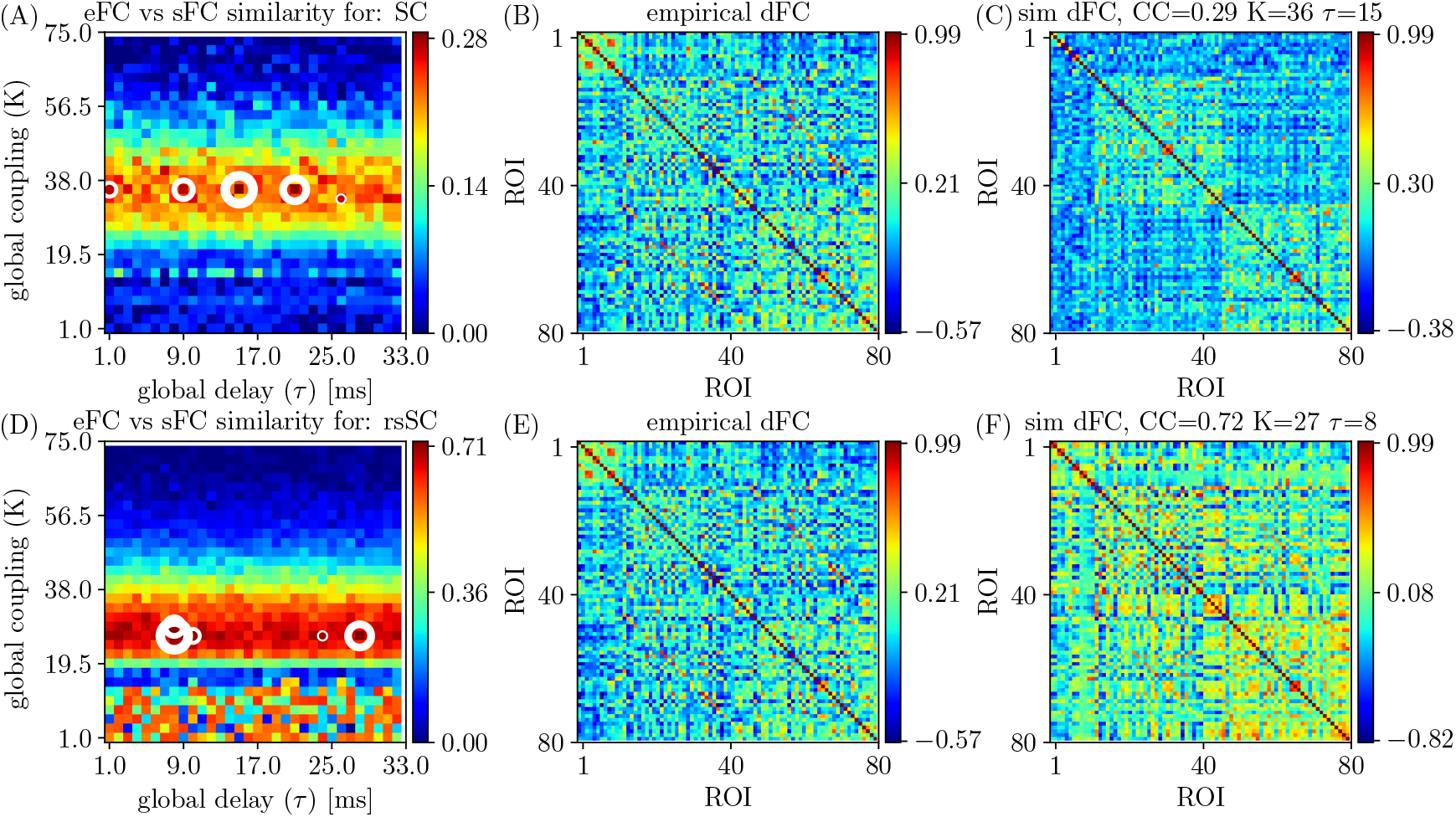
Parameter Sweep Exploration for Phase Coherence Connectivity (edFC vs. sdFC). Upper row (using the respective subject’s standard SC matrix to define the weights in model Eq. (8): (A) The colormap depicts the CC_dFC_ = corr(sdFC,edFC) for the parameters (*K, τ*). (B) The edFC calculated from the empirical BOLD signal. (C) The sFC matrix with the larger CC_dFC_. Lower row (using the respective subject’s hybrid rsSC matrix to define the weights in model Eq. (8): (D) The respective colormap for CC_dFC_ = corr(sdFC,edFC). (E) The edFC calculated from the empirical BOLD signal (same as (B)). (E) The sdFC matrix with the larger CC_dFC_. Note the different ranges in the respective colorbars of panels (B), (C), (E) and (F) capturing the different temporal alignment of the phase of the respective BOLD signals. See text for more details.

The optimal match between sdFC and edFC in the parameter space is acquired for (*K, τ*)-values where CC_dFC_ becomes maximal (in panels (A) and (D) we indicate 5 maximum values with white circles. The upper row shows the results when we use the respective subject’s standard SC matrix to define the weights in model Eq. (8). **Figure** 7(A) depicts the CC_dFC_ = corr(sdFC,edFC) for the parameters (*K, τ*), while **Figure** 7(B) the edFC calculated from the empirical BOLD signal. **Figure** 7(C) shows the sdFC matrix obtained by the larger CC_dFC_. In the lower row, we show the same analysis using now the respective subject’s hybrid rsSC matrix. Once again, we find that the use of hybrid rsSC yields substantial improvement in the best fit between empirical and simulated BOLD activity (CC_dFC_(rsSC) ≈ 0.72 while CC_dFC_(SC) ≈ 0.29), this time in the context of dynamical functional connectivity (in the Supplementary Materials **Figure** A.1. we present the corresponding correlation analysis and scatterplots). Here, we have presented the output for the same example subject (as the one in previous figures). However, this conclusion holds for all subjects similar to what we did in **Figure** 4 (in the Supplementary Materials **Figure** A.2. we show the respective statistical analysis and boxplots).

## 4. Discussion

In this study, we showed that a coupled Kuramoto oscillator system built on a novel brain connectome can yield simulated BOLD brain activities that strongly resemble actual BOLD signals observed during resting-state fMRI. This novel brain connectome rsSC combines characteristics of both structural and functional information, such that the sign of an edge indicates whether the coupling between two brain regions is excitatory (i.e., attractive) or inhibitory (i.e., repulsive). We used the TVB computational platform with the Kuramoto model [57] and generated simulated BOLD time series across a range of different model parameters (*K, τ*) (producing PSE colormaps like in **Figure** 2). This allowed us to optimize model parameters and tune generated synthetic BOLD signals that produce simulated functional connectivity (FC) most similar to actual observed FC.

Our testing sample is an APOE *ε*4 dataset [60] that contains both structural (traditional SC and rsSC) and functional (BOLD signals and FC matrices) for two groups of cognitively-normal subjects, namely Carriers and Non Carriers (control). Our ultimate goal was to numerically compare the performance of Kuramoto oscillator models in producing simulated FC when using traditional SC versus rsSC (see e.g. [18, 40, 61, 21]) which define the pair-wise coupling weights (*c*_*ij*_) in the Kuramoto model Eq. (8) in fundamentally different ways (see **Figure** 1), with traditional SC only permits attractive coupling. As the tract lengths were not available, we considered the euclidean distances between ROIs in the Desikan atlas [59] to account for the mean delay parameter (*τ*) in the Kuramoto model.

Overall, we found that there are important advantages in using hybrid rsSC as it can produce BOLD sequences and synthetic FC that follow well the general trends of the empirical BOLD time series and empirical FC (**Figures** 2 and 4). Indeed, hybrid rsSC performed significantly better than standard SC matrices as measured by their correlation values CC_FC_ = corr(sFC,eFC) (scatterplots in **Figure** 3). The use of standard SC matrices have substantially lower CC_FC_ than when using rsSC instead (see colorbars in **Figure** 2(A) and (D)). The statistical analysis and overall better performance of rsSC is presented in **Figure** 4 summarizing our finding for all (76 in total) subjects, showing the striking superiority of the rsSC over traditional SC in generating simulated BOLD signals (see **Figure** 5). A similar conclusion was also reached when we repeated PSE analysis using dynamic instead of static Functional Connectivity (**Figure** 7) matrices.

As the hybrid rsSC and traditional SC differ the most in the former’s ability to capture inhibitory coupling, we further examine the simulation performance w.r.t. positive versus negative BOLD correlations. Despite the fact that in general both sFC (simulated with rsSC/SC matrices) perform rather well in capturing the positive correlations observed in the empirical BOLD signals, only the rsSC ones can effectively produce negative correlations closely matching those occurring in the empirical BOLD signals (see also [75]). This result was demonstrated in **Figure** 6 and was observed for both Non Carriers and Carriers datasets.

Our study has a few limitations. First, we restricted ourselves to a specific frequency band during simulations and thus future studies should further explore different ranges of frequencies in the Kuramoto model, e.g. either in different Hz ranges or extracted directly from the empirical BOLD signals per node and per subject (see e.g [63, 16] and references therein). Furthermore one may validate these findings for different dynamical models, or to furthermore consider additional relevant dynamical features such as noise or the use of neuroimaging data where the path lengths is also available. Let us also stress that in this work we do not seek to detect model parameters settings that could distinguish between Carriers and Non Carriers based on the presence or not of the APOE *ε*4 gene or age and gender factors.

In summary, here we showed that our recently proposed hybrid connectome rsSC can produce simulated synthetic BOLD signals that yield functional connectivity matrices strikingly similar to those actually obtained during the resting-state. The construction of rsSC and subsequent Kuramoto oscillator based simulations can be implemented in a relatively straightforward fashion, requiring only the tractography-derived (weights and tract lengths) structural and resting-state functional connectome from a standard neuroimaging preprocessing pipeline. Thus, we conclude by highlighting that existing publicly available open-source pipelines, such as the TVB platform, could be easily equipped to include an add-on module that incorporates rsSC for the neuroscientific community interested in the modeling of simulated fMRI BOLD time series.

## Acknowledgments

We acknowledge the use of Fenix Infrastructure resources, which are partially funded from the European Union’s Horizon 2020 research and innovation programme through the ICEI project under the grant agreement No. 800858. In particular, we acknowledge the access to the JUSUF supercomputer at the Jülich Supercomputer Centre. TM would like to thank the Institut Henri Poincaré for its support through the ‘Research in Paris’ programme and hospitality in the period during which part of this work took place.

## Data and code availability statement

The original contributions presented in the study are included in the article, further inquiries can be directed to the corresponding author. The TVB code used for the simulation of the brain network dynamics can be found in this repository: https://gitlab.jsc.fz-juelich.de/metaopt/rssc_simulations_tvb.git. The scripts for generating rsSC are available here: https://github.com/iforte2/hybrid-connectome. Regarding the statistical tests used in the manuscript, standard python scripts were used (written by the authors).

## Ethics statement

This study represents a secondary data analysis using imaging data obtained as part of a larger study. See Korthauer et al. [60] for additional information on the study and the relevant institutional approval.

## CRediT authorship contribution statement

**Thanos Manos**: Conceptualization, Methodology, Software, Resources, Formal analysis, Writing - original draft, review and editing, Visualization, Supervision, Project administration, Funding acquisition. **Sandra Diaz-Pier**: Conceptualization, Methodology, Software, Resources, Writing review and editing. **Igor Fortel**: Software, Data curation, Writing review and editing. **Ira Driscoll**: Software, Formal analysis, Data curation. **Liang Zhan**: Software, Data curation, Writing review and editing. **Alex Leow**: Conceptualization, Methodology, Software, Formal analysis, Writing review and editing, Supervision, Funding acquisition.

## Appendix A. Supplementary materials

**Figure 8:**
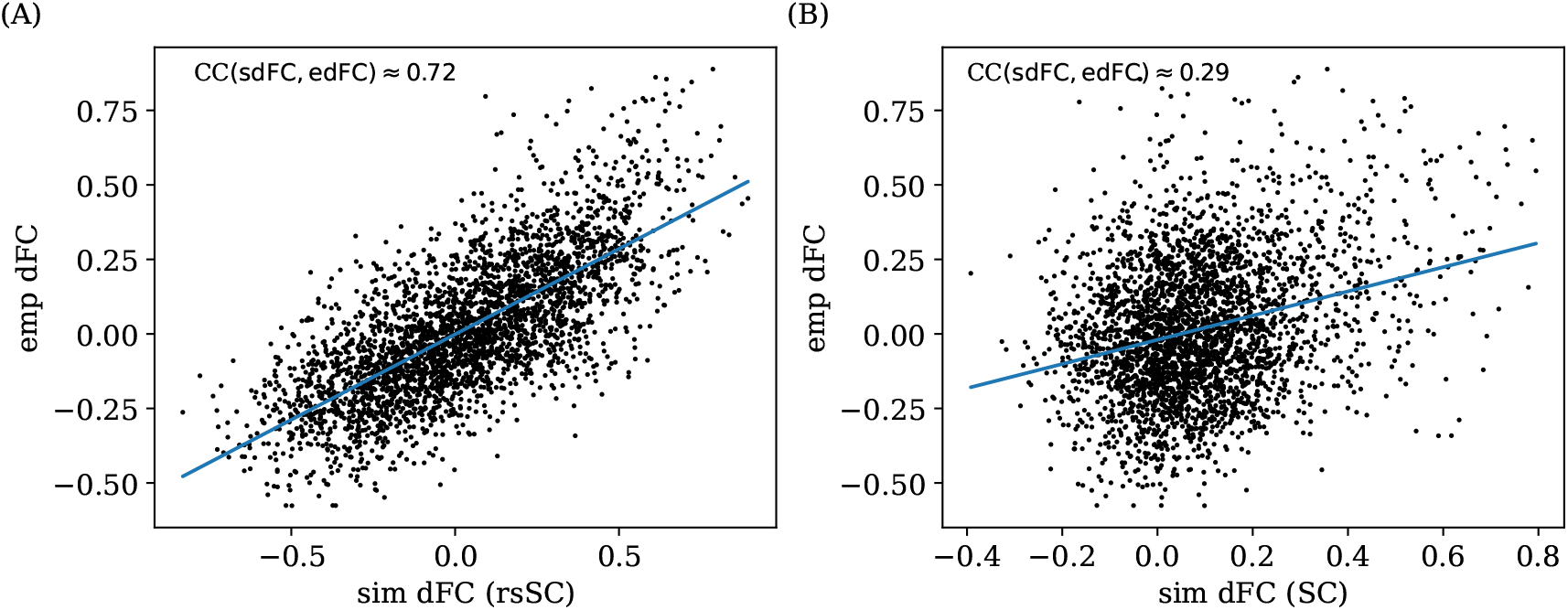
Correlation Analysis. Scatterplots between empirical (*y*−axis) and optimal simulated dFC correlations (*x*−axis) aggregated across all entries in the corresponding dFC matrices, i.e., panels (B) and (C) in **Figure** 7. Note that a perfect match between the two dFC matrices would thus place all the points along the line *x* = *y*. (A) edFC vs. sdFC using rsSC (B) edFC vs. sdFC using standard SC. Both panels refer to same subject presented in **Figure 7**). The blue lines indicate the respective linear regression model.

**Figure 9:**
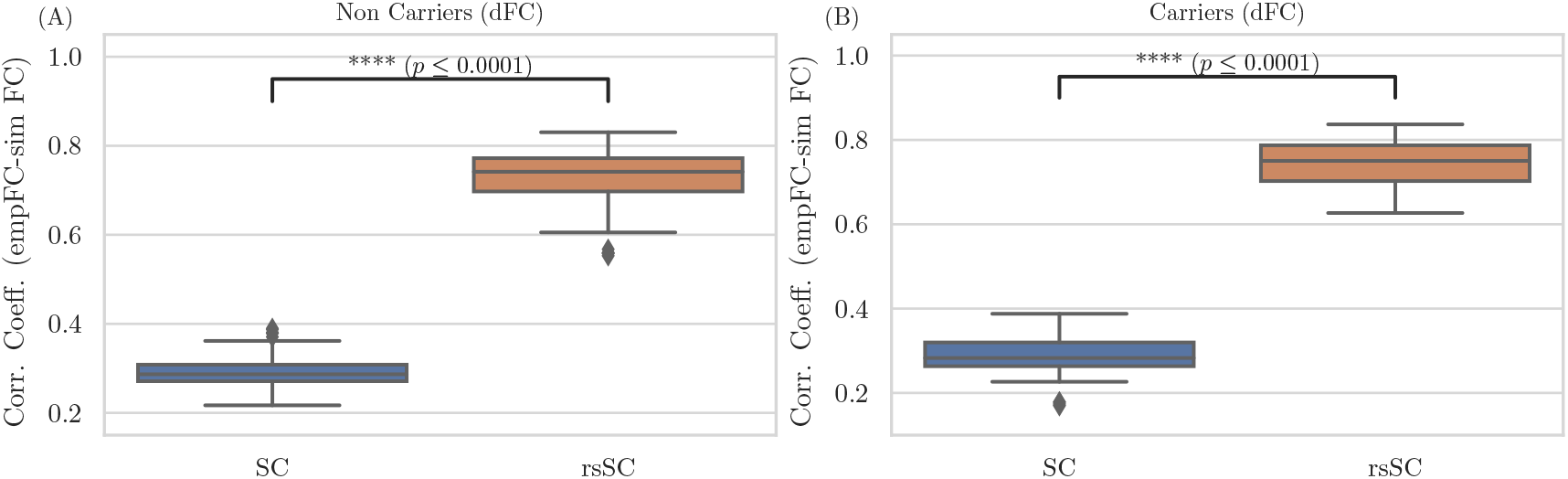
Statistical Analysis. Boxplots for the non-carriers (A) and carriers (B) dataset (38 subjects per type) correlation coefficients between edFC and sdFC using SC and rsSC matrices for the simulated time series respectively. For each subject we considered the 5 maximum values (see circles in **Figures 7**(A),(D)). The difference in the respective mean values of the two datasets is statistically significant measured by the t-test with very small *p*−value (*p* ≤ 0.0001) for both non-carriers and carriers sets.

